# A lateralized sensory signaling pathway mediates context-dependent olfactory plasticity in *C. elegans*

**DOI:** 10.1101/2025.07.25.666858

**Authors:** Anjali Pandey, Maya Katz, Stephen Nurrish, Alison Philbrook, Piali Sengupta

**Affiliations:** Department of Biology, Brandeis University, Waltham, MA 02454; Department of Biology, Rhode Island College, Providence, RI 02908

**Keywords:** olfaction, plasticity, AWC, left-right asymmetry, *C. elegans*, guanylyl cyclase

## Abstract

Lateralization of neuronal functions plays a critical role in regulating behavioral flexibility, but the underlying molecular mechanisms are challenging to establish at a single-neuron level. We previously showed that attraction of *C. elegans* to a medium-chain alcohol switches to avoidance in a uniform background of a second attractive odorant. This context-dependent behavioral plasticity is mediated by symmetric inversion of the odor-evoked response sign in the bilateral AWC olfactory neurons. Here we show that this symmetric response plasticity is driven by asymmetric molecular mechanisms in the AWC neuron pair. Mutations in the *gcy-12* receptor guanylyl cyclase abolish odor response plasticity only in AWC^OFF^; the opposing odor-evoked response signs in AWC^OFF^ and AWC^ON^ in *gcy-12* mutants results in these animals being behaviorally indifferent to this chemical. We find that *gcy-12* is expressed, and required, in both AWC neurons to regulate odor response plasticity only in AWC^OFF^. We further show that disruption of AWC fate lateralization results in loss of asymmetry in the response plasticity in *gcy-12* mutants. Our results indicate that symmetric neuronal response plasticity can arise from asymmetry in underlying molecular mechanisms, and suggest that lateralization of signaling pathways in defined conditions may enhance neuronal and behavioral flexibility.

## INTRODUCTION

Animals continuously sample their complex chemical environments to inform their behavioral and developmental decisions. While subsets of chemicals can be innately attractive or aversive, the ability to modify these responses based on experience, context, and internal state enables organisms to adapt their behaviors for optimal survival and reproduction [1–3]. Decades of work have described neuronal mechanisms that underlie the generation of chemosensory behavioral plasticity. In addition to extensively characterized pathways that operate centrally to integrate chemosensory stimuli with other inputs [2, 4–6], altered responses of peripheral chemosensory neurons can also contribute to behavioral flexibility (eg. [7–9]). The molecular pathways regulating response plasticity in defined individual chemosensory neuron types remain to be fully described.

Left-right asymmetry of sensory inputs and processing (lateralization) is a key characteristic of the brain across species and is thought to in part expand the brain’s remarkable cognitive capacities [10–14]. Lateralization is critical for depth perception, localization of a sound source, and tracking odor gradients among other behaviors [15–19]. In principle, sensory lateralization can arise from asymmetry at any stage in the neural circuit including at the level of peripheral sensory neuron responses, although the complexity of most animal nervous systems makes it challenging to provide a detailed analysis of functional asymmetry at the level of individual neuron types.

*C. elegans* detects and discriminates among hundreds of chemicals using a small, well-characterized set of chemosensory neurons, a subset of which is present in the bilateral amphid sensory organs of the head [20–22]. The ASE and AWC amphid sensory neuron pairs exhibit left-right asymmetry in both their transcriptional profiles and response properties [23–29]. While the left-right asymmetric ASE fates are developmentally specified, the fates of the two AWC olfactory neurons are determined stochastically via calcium signaling during early development [29–31]. Thus, either the left or right AWC neuron assumes the ‘AWC^ON^’ fate as defined by the expression of the *str-2* chemoreceptor, whereas the other expresses the ‘AWC^OFF^’ fate and expresses a partly distinct set of chemoreceptors including *srsx-3* but not *str-2* [29, 31–33]. This left-right sensory neuronal asymmetry is thought to contribute to the diversification and expansion of this organism’s chemosensory repertoire and aid in odorant discrimination.

Similar to other animals, olfactory behaviors in *C. elegans* are extensively modulated in an experience- and state-dependent manner [2]. In addition to central mechanisms, modulation of primary responses in chemosensory neurons plays a major role in driving behavioral plasticity in *C. elegans* (eg. [7, 34–36]). The known functions and molecular profiles of each chemosensory neuron type, together with the ability to monitor stimulus-evoked neuronal responses in living animals [21, 33], provide an opportunity to establish the mechanisms by which plasticity in defined sensory neuron response properties can drive behavioral flexibility.

AWC senses multiple bacterially-produced volatile attractive odorants including low concentrations of the alcohols 1-hexanol (henceforth referred to as hexanol) and isoamyl alcohol (IAA) [20, 37]. Addition and removal of these chemicals decreases and increases intracellular calcium concentrations, respectively, in both AWC neurons thereby driving attraction [37–39]. We previously showed that in low uniform concentrations of IAA (saturating IAA: sIAA), hexanol instead activates both AWC neurons resulting in animals strongly avoiding hexanol [37] (Fig 1A). The ODR-1 receptor guanylyl cyclase is necessary for attraction to hexanol, whereas the ODR-3 Gα subunit is required for avoidance of hexanol in sIAA conditions [37] (Fig 1A). Additional molecules and mechanisms mediating this context-dependent odorant response plasticity remain to be identified.

**Fig 1.**
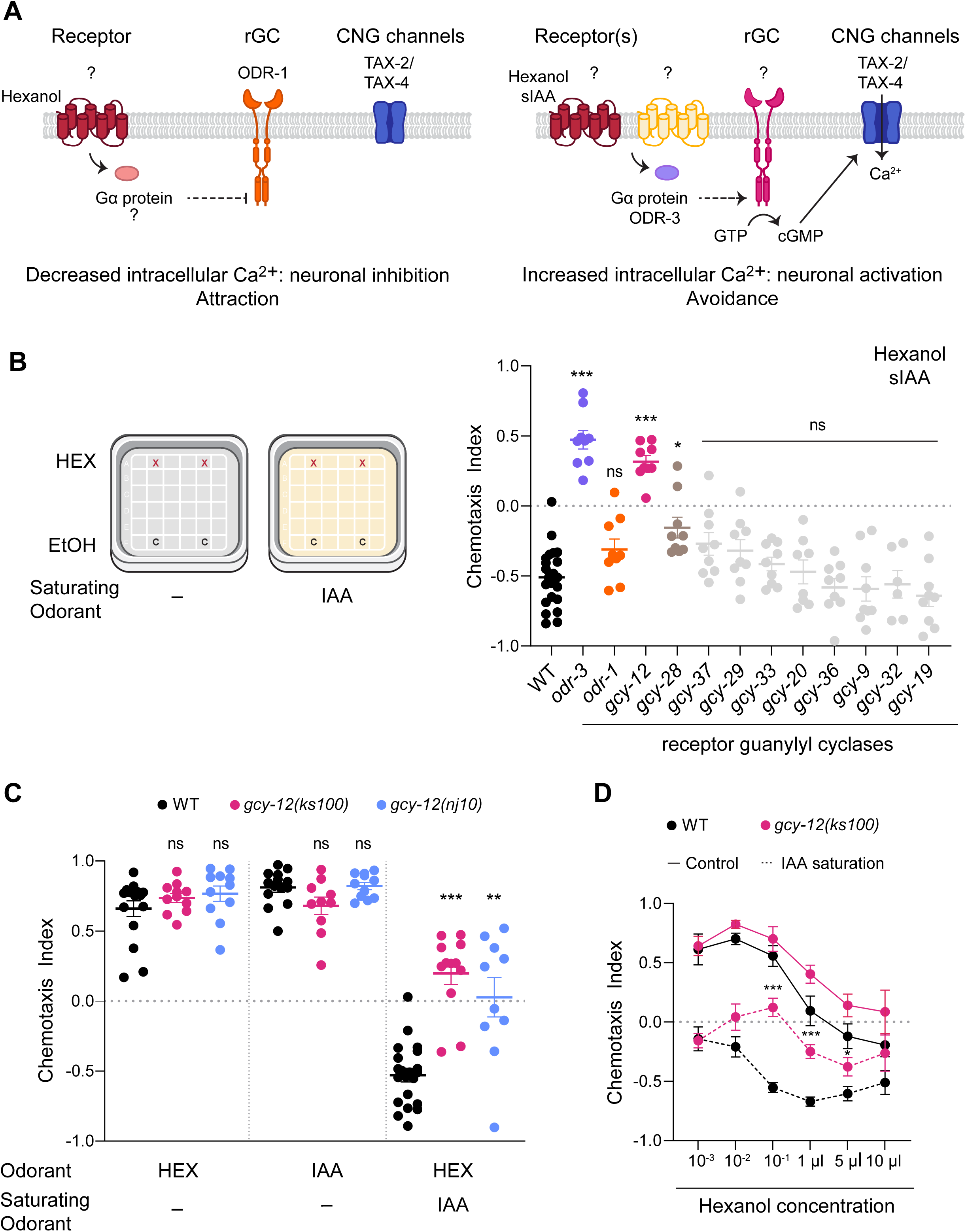
*gcy-12* mutants exhibit defects in hexanol avoidance in sIAA. **A)** Odorants such as hexanol decrease intracellular calcium and inhibit AWC to drive attraction. In sIAA, hexanol instead increases intracellular calcium and activates AWC to promote avoidance. The ODR-1 receptor guanylyl cyclase and ODR-3 Gα subunit are required for hexanol attraction in control conditions, and avoidance in sIAA, respectively [37]. **B)** (Left) Cartoon of the control and sIAA conditions for chemotaxis behavioral assays. (Right) Behavioral responses to a point source of 10^-1^ hexanol (HEX) in 10^-4^ sIAA. Positive and negative chemotaxis index values indicate attraction and avoidance, respectively. Alleles are listed in S1 Table. Each dot is the chemotaxis index calculated from a single assay of ∼50-100 animals. Data shown are from at least three independent days. Horizontal and vertical bars are the mean and SEM, respectively. * and ***: different from wild-type at P<0.05 and P<0.001, respectively (Kruskal-Wallis with Dunn’s post-hoc correction). ns: not significant. **C)** Behavioral responses of wild-type and *gcy-12* mutants to the shown odorants in the absence or presence of sIAA. Concentrations of odorants used were 10^-1^ hexanol or IAA as test odorants, and 10^-4^ IAA as the saturating odorant. Each dot is the chemotaxis index calculated from a single assay of ∼50-100 animals. Data shown are from at least three independent days. Horizontal and vertical bars are the mean and SEM, respectively. ** and ***: different from corresponding wild-type at P<0.01 and 0.001, respectively (Kruskal-Wallis with Dunn’s post-hoc correction). ns: not significant. **D)** Responses of wild-type and *gcy-12(ks100)* mutants to the indicated concentrations of hexanol in 10^-4^ sIAA. Each filled circle is the average value from six chemotaxis behavioral assays of ∼50-100 animals performed over three days. Errors are SEM. * and ***: different from corresponding wild-type at P<0.05 and 0.001, respectively (Kruskal-Wallis with Dunn’s post-hoc correction).

Here we describe a role for the GCY-12 receptor guanylyl cyclase in mediating left-right asymmetric context-dependent plasticity in hexanol responses in AWC. While AWC^ON^ is activated upon the addition of hexanol in both wild-type and *gcy-12* mutants in sIAA, AWC^OFF^ is instead inhibited upon addition and activated upon removal of hexanol in sIAA in *gcy-12* mutants. The asymmetric and opposing hexanol-evoked responses in AWC^OFF^ and AWC^ON^ result in *gcy-12* mutants being behaviorally indifferent to hexanol in sIAA. *gcy-12* is expressed, and required, in both AWC neurons to mediate asymmetric hexanol response plasticity defects. We further find that mutations that disrupt AWC fate lateralization also abolish the asymmetric response plasticity phenotype of *gcy-12* mutants. Our results uncover cryptic asymmetry in the mechanisms underlying symmetric response plasticity in an olfactory neuron type, and suggest that asymmetric context-dependent reconfiguration of the signal transduction pathway in sensory neurons provides additional mechanistic flexibility in generating behavioral plasticity.

## RESULTS

### The GCY-12 receptor guanylyl cyclase is necessary for the context-dependent switch in hexanol response valence

Mutations in the *tax-4* cyclic nucleotide-gated channel subunit abolish both attraction to, and avoidance of, hexanol in control (i.e. without saturating odorants) and sIAA conditions, respectively [37], indicating that cGMP-mediated signaling underlies both behaviors (Fig 1A). We previously showed that the ODR-1 receptor guanylyl cyclase is necessary to drive attraction to hexanol in control conditions, but has no effect on hexanol response plasticity in sIAA [37] (Fig 1A). These observations suggest that additional guanylyl cyclase(s) likely operate to invert the response sign in AWC and drive hexanol avoidance in sIAA.

Multiple receptor and soluble guanylyl cyclases including ODR-1 are predicted to be expressed in the two AWC neurons [23, 25, 33, 40, 41], with a subset exhibiting possible asymmetric expression levels in AWC^ON^ and AWC^OFF^ (S1A Fig). As we showed previously [37], *odr-1* mutants avoided hexanol in sIAA similar to wild-type animals, whereas animals mutant for the *odr-3* Gα subunit failed to avoid this odorant and were instead strongly attracted (Fig 1B). We found that *gcy-12* and *gcy-28* mutants also failed to robustly avoid hexanol in sIAA; these mutants were instead more weakly attracted, or indifferent, to hexanol (Fig 1B). GCY-28 localizes to AWC^ON^ axons and regulates olfactory behavior via modulation of synaptic signaling but not primary olfactory signal transduction [42]. GCY-12 is expressed broadly in multiple sensory neuron types including in AWC [33, 43], and has been implicated in the regulation of body size and neuronal gene expression [43, 44]. Although *gcy-12* mutants have not been reported to exhibit olfactory behavioral defects [43], a GCY-12::GFP fusion protein localizes to the cilia or distal dendritic ends of AWC and other sensory neurons [43] (see further below). We thus focused further on a possible role of GCY-12 in driving context-dependent plasticity in hexanol responses in AWC.

The *gcy-12(ks100)* nonsense and *gcy-12(nj10)* deletion alleles (S1B Fig) are predicted to encode truncated proteins lacking the intracellular catalytic domain [43]. Both mutants were either indifferent to or exhibited weak attraction to hexanol in sIAA, although their responses to hexanol and IAA alone under control conditions were unaffected (Fig 1C). Moreover, *gcy-12* mutants exhibited reduced attraction to a point source of IAA in sIAA similar to wild-type animals (compare attraction to IAA alone in Fig 1B to IAA in sIAA in S1C Fig), suggesting that the inability of *gcy-12* mutants to avoid hexanol in sIAA is unlikely to arise simply due to defects in IAA detection or saturation.

Wild-type animals are attracted to low hexanol concentrations but avoid high concentrations [37] (Fig 1D). This attraction and avoidance are mediated by the AWC olfactory and ASH nociceptive neurons, respectively [37]. In sIAA, wild-type animals instead avoid hexanol at all concentrations [37] (Fig 1D). Avoidance of low hexanol concentrations in sIAA is mediated via inversion of the hexanol-evoked calcium response in AWC [37] (Fig 1A). While wild-type and *gcy-12* mutants exhibited similar behaviors to a range of hexanol concentrations under control conditions, *gcy-12* mutants were indifferent to hexanol at low concentrations but retained the ability to avoid high hexanol concentrations in sIAA (Fig 1D). These results suggest that GCY-12 may specifically regulate the plasticity of hexanol responses in AWC in sIAA conditions.

### GCY-12 regulates context-dependent inversion in hexanol responses only in the AWC^OFF^ neuron

To determine how mutations in *gcy-12* affect hexanol responses, we examined stimulus-evoked intracellular calcium dynamics in the bilateral AWC neurons. Consistent with previous work [37], nearly all AWC neurons in wild-type animals responded to hexanol in sIAA by increasing intracellular calcium levels (S2A Fig). However, we noted that only ∼50% of AWC neurons in *gcy-12* mutants exhibited the expected sign inversion of the hexanol response in sIAA (S2A Fig). Since these neurons were imaged without a bias towards AWC^ON^ or AWC^OFF^, these observations raised the possibility that mutations in *gcy-12* affect hexanol responses in AWC in sIAA in an asymmetric manner.

We addressed this notion by comparing hexanol-evoked calcium responses specifically in AWC^ON^ or AWC^OFF^ neurons; AWC^OFF^ was identified via expression of an *srsx-3*p::mScarlet reporter [32] (see Materials and Methods). Addition of either hexanol or IAA alone altered intracellular calcium dynamics in the soma of both AWC neurons similarly with relatively minor differences in response amplitude and dynamics between wild-type and *gcy-12* mutants (Fig 2A and 2B). IAA-evoked responses were also reduced in sIAA in both AWC neurons in *gcy-12* mutants (S2B Fig) consistent with the reduced attraction of these animals to a point source of IAA under these conditions (S1C Fig).

**Fig 2.**
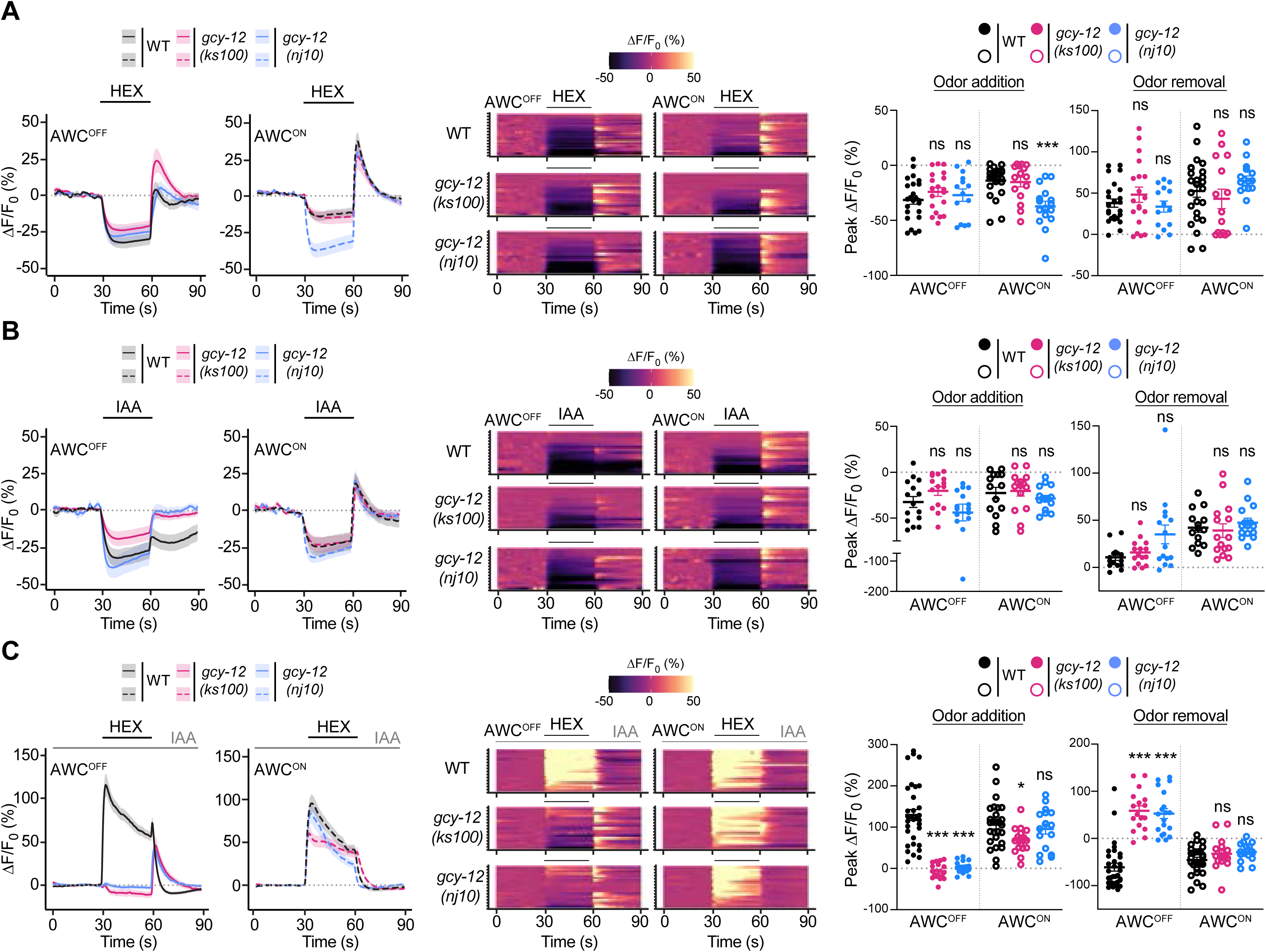
*gcy-12* mutants exhibit defects in context-dependent hexanol response plasticity only in AWC^OFF^. **A-C)** (Left) Average changes in GCaMP fluorescence in AWC^OFF^ and AWC^ON^ in wild-type and *gcy-12* mutants upon addition of a 30 sec pulse of 10^-4^ hexanol (A,C) or IAA (B) indicated by a short solid line. The presence of a 10^-4^ concentration of sIAA in C is indicated by a long solid gray line. Shaded regions are SEM. (Middle) Heatmaps of changes in fluorescence intensity corresponding to the responses shown at left in each panel. Each row in the heatmaps shows responses from a single AWC^OFF^ or AWC^ON^ neuron from different animals. (Right) Quantification of peak changes in fluorescence intensity in AWC^OFF^ or AWC^ON^ upon odorant addition or removal in each panel. Each circle is the value from a single neuron. Horizontal and vertical bars indicate the mean and SEM, respectively. * and ***: different from corresponding wild-type at P<0.05 and 0.001, respectively (Kruskal-Wallis with Dunn’s posthoc correction). ns: not significant.

As we reported previously, hexanol robustly increased intracellular calcium levels in both AWC neurons in sIAA in wild-type animals (Fig 2C, S1 and S2 Videos) [37]. However, while hexanol also increased calcium levels in AWC^ON^ in *gcy-12* mutants in sIAA, addition of hexanol instead weakly decreased, and removal strongly increased, intracellular calcium levels in sIAA in AWC^OFF^ in these animals (Fig 2C, S1 and S2 Videos). Hexanol-evoked calcium responses in the ASH neurons were unaffected upon loss of *gcy-12* (S2C Fig). Hexanol response defects were also observed asymmetrically only in the AWC^OFF^ neuronal cilia in sIAA but not under control conditions in *gcy-12* mutants (S3 Fig). We infer that GCY-12 is necessary for the hexanol-evoked inversion in the sign of the calcium response specifically in AWC^OFF^ in sIAA but is dispensable for odorant responses under control conditions. Since AWC-mediated attraction and avoidance behaviors are correlated with stimulus-evoked decreases and increases in calcium levels in AWC, respectively [37, 38, 45], the conflicting asymmetric responses in AWC^OFF^ and AWC^ON^ upon hexanol addition and removal in *gcy-12* mutants likely account for the behavioral indifference of these animals to hexanol in sIAA.

### Olfactory plasticity defects in *gcy-12* mutants are partly odorant-specific

We previously showed that the sIAA-dependent response inversion in AWC is not restricted to hexanol, but that the response to the typically attractive odorant 1-heptanol (henceforth referred to as heptanol) is also inverted in sIAA resulting in avoidance of this chemical [37]. Moreover, saturation with a second AWC-sensed attractive chemical benzaldehyde also results in inversion of the response to hexanol in AWC [37]. We tested whether mutations in *gcy-12* affect context-dependent odorant response plasticity more broadly in AWC, and whether these effects are asymmetric.

Both wild-type and *gcy-12* mutants were attracted to heptanol under control conditions (Fig 3A). Consistently, heptanol-evoked calcium responses were also similar in both AWC neurons in wild-type and *gcy-12* mutants in control conditions (Fig 3B, S4A Fig). Although *gcy-12* mutants retained the ability to avoid heptanol in sIAA, this behavior was weakly but significantly reduced in these mutants as compared to the behaviors of wild-type animals (Fig 3A). Heptanol activated both AWC neurons in wild-type and *gcy-12* mutants in sIAA (Fig 3C, S4B Fig), likely accounting for the ability of these animals to avoid heptanol. However, in *gcy-12* mutants, AWC^OFF^ consistently exhibited weaker activation upon heptanol addition, and stronger activation upon heptanol removal, as compared to wild-type animals in sIAA (Fig 3C, S4B Fig), suggesting that GCY-12 may also partly modulate heptanol responses asymmetrically in AWC^OFF^ in sIAA to regulate behavioral plasticity.

**Fig 3.**
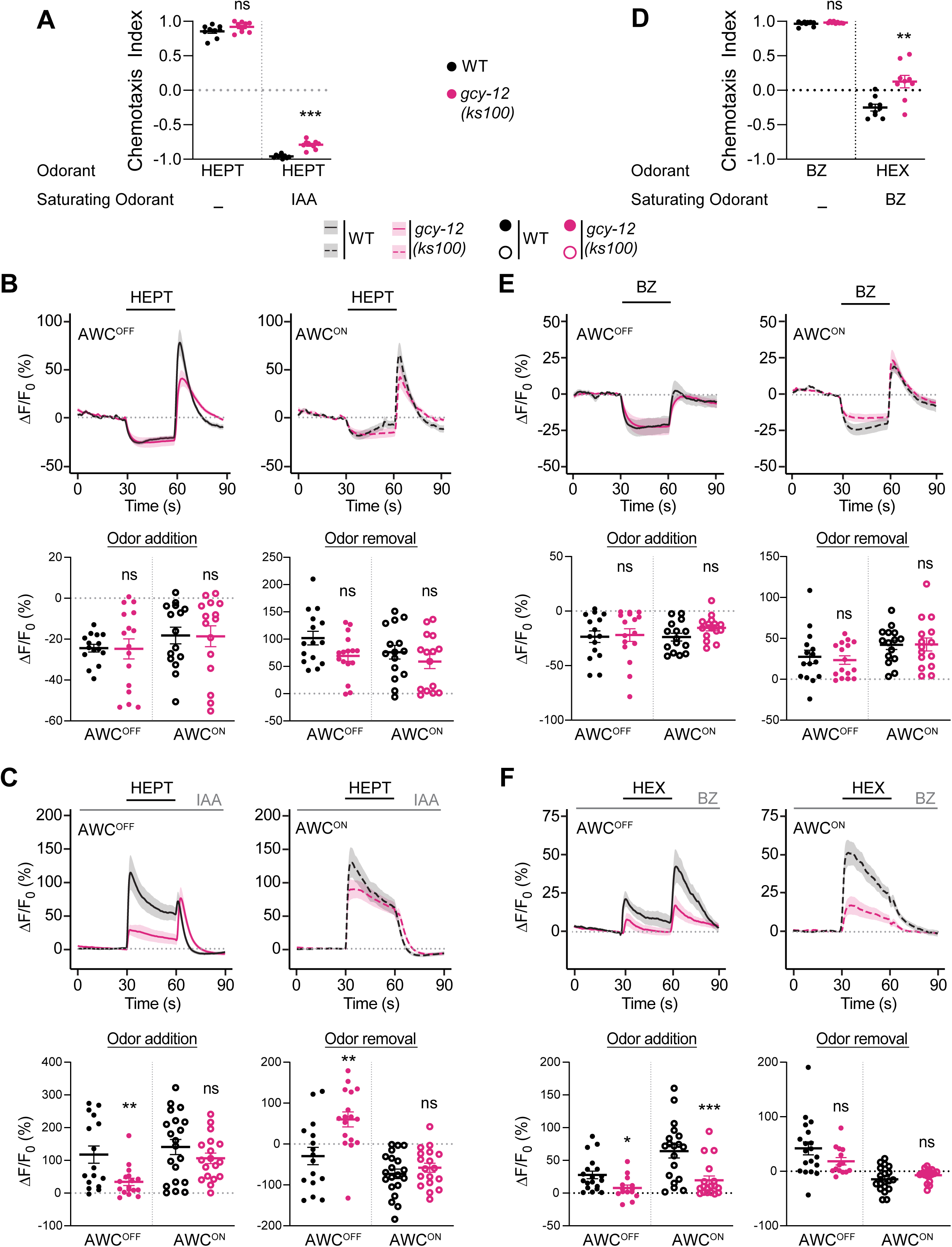
Context-dependent odorant response plasticity in *gcy-12* mutants may be partly odorant-specific. **A, D)** Behavioral responses of wild-type and *gcy-12* mutants to point sources of the indicated odorants in sIAA (A) or saturating benzaldehyde (D: BZ). Concentrations of odorants used were 10^-1^ heptanol or hexanol, and 1:200 dilution of benzaldehyde as test odorants, and 10^-4^ IAA or benzaldehyde as the saturating odorants. Each dot is the chemotaxis index calculated from a single assay of ∼50-100 animals. Data shown are from at least three independent days. Horizontal and vertical bars are the mean and SEM, respectively. ** and ***: different from corresponding wild-type at P<0.01 and 0.001, respectively (A: unpaired t-test with a post-hoc Welch’s correction; D: Mann-Whitney-Wilcoxon test).). ns: not significant. **B, C, E, F)** (Top) Average changes in GCaMP fluorescence in AWC^OFF^ and AWC^ON^ in wild-type and *gcy-12* mutants upon addition of a 30 sec pulse of 10^-4^ heptanol (B, C), benzaldehyde (E) or hexanol (F) indicated by a short solid line. The presence of a 10^-4^ concentration of sIAA (C) or saturating benzaldehyde (F) is indicated by a long solid gray line. Shaded regions are SEM. (Bottom) Quantification of peak changes in fluorescence intensity in AWC^OFF^ or AWC^ON^ upon odorant addition or removal in each panel, corresponding to the responses shown at top. Each circle is the value from a single neuron. Corresponding heatmaps are shown in S4 Fig. Horizontal and vertical bars indicate the mean and SEM, respectively. *, **, and ***: different from corresponding wild-type at P<0.05, <0.01, and 0.001, respectively (Mann-Whitney-Wilcoxon test). ns: not significant.

In wild-type animals, attraction to hexanol switched to weak aversion in saturating concentrations of benzaldehyde, although *gcy-12* mutants were largely indifferent to hexanol under these conditions (Fig 3D). Benzaldehyde elicited similar symmetric responses in both AWC neurons in wild-type and *gcy-12* mutants (Fig 3E, S4C Fig). However, hexanol-evoked calcium responses in saturating benzaldehyde were asymmetric even in wild-type animals (Fig 3F, S4D Fig). Response amplitudes but not response signs were altered upon hexanol addition in saturating benzaldehyde in both AWC neurons in *gcy-12* mutants (Fig 3F, S4D Fig). We conclude that while GCY-12 regulates hexanol and heptanol response plasticity asymmetrically in AWC^OFF^ in sIAA, this molecule may also contribute to the modulation of olfactory response plasticity in both AWC neurons in other odorant contexts.

### Symmetric loss of *gcy-12* is necessary for asymmetric odorant response plasticity defects

Neuronal transcriptomics data indicate that *gcy-12* is expressed in multiple neuron types including at very low levels in both AWC neurons with expression levels in AWC^OFF^ reported to be twice that in AWC^ON^ (S1A Fig) [33]. Overexpression of a genomic *gcy-12::gfp* transgene also showed expression in multiple neurons including AWC [43]. The locus is predicted to encode two protein isoforms containing all predicted extracellular and intracellular domains including the catalytic domain, with the longer GCY-12.a protein containing additional C-terminal sequences (S1B Fig). Analyses of neuron type-specific alternative splicing data indicate that 90% and 10% of transcripts in AWC encode GCY-12.a and GCY-12.b, respectively [46]. To assess the expression pattern and subcellular localization of individual isoforms, we engineered an endogenous allele predicted to encode reporter-tagged GCY-12.a and GCY-12.b isoforms (S5A Fig). Consistent with low expression in AWC reported by transcriptional profiling [33], we detected very weak expression of endogenously tagged GCY-12.a in a subset of neuronal soma in the head; expression of endogenously tagged GCY-12.b was undetectable (S5B Fig). Expression of the *gcy-12.a* fusion gene was too weak to definitively identify individual neurons or to establish subcellular localization of the encoded protein.

Overexpressed GCY-12.b was shown to be enriched in the proximal regions of sensory cilia of AWC and additional sensory neurons [43]. Since endogenous expression levels were too low to allow assessment of protein localization, we determined the subcellular localization of each isoform by expressing fluorescent reporter-tagged *gcy-12.a* and *gcy-12.b* cDNAs under the *odr-1* promoter that drives expression in both AWC neurons [40]. Both reporter-tagged isoforms were present in the soma, dendrites and AWC sensory endings (Fig 4A) although we did not observe localization to axons. As reported previously [43], expression of these fusion proteins was punctate in the dendrites and at the AWC sensory endings (Fig 4A). We infer that GCY-12 can localize to the sensory cilia of both AWC neurons.

**Fig 4.**
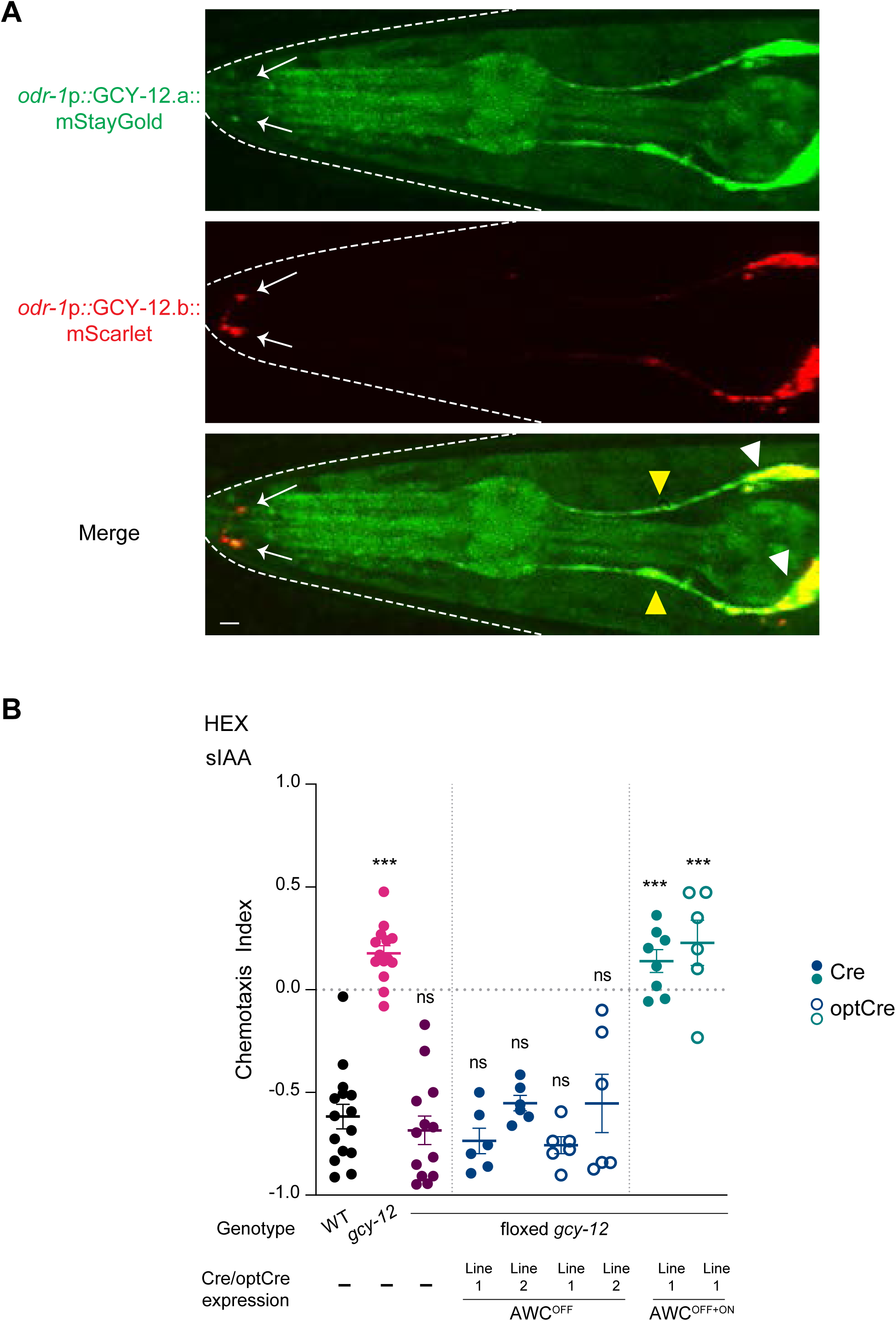
GCY-12 is localized to the cilia of both AWC neurons and acts in both neurons to regulate hexanol response plasticity in AWC^OFF^. **A)** Representative images of adult hermaphrodites expressing *odr-1*p*::gcy-12.a::mStayGold* and *odr-1*p*::gcy-12.b::mScarlet*. Arrows point to the AWC sensory endings. White and yellow arrowheads indicate soma and dendrites, respectively. The worm head is outlined by dashed lines. Anterior is at left. Scale bar: 5 μm. **B)** Behavioral responses of animals of the indicated genotypes to a point source of 10^-4^ hexanol in sIAA. Cre was expressed in AWC^OFF^ or both AWC neurons under the *srsx-3* (AWC^OFF^) or *odr-1*(AWC^OFF+ON^) promoters, respectively. Cre function was optimized (optCre) by replacing the SV40 with the strong NLS from *egl-13* and injected at a higher concentration (see Materials and Methods) [67, 68]. Each dot is the chemotaxis index calculated from a single assay of ∼50-100 animals. Data shown are from at least three independent days. Horizontal and vertical bars are the mean and SEM, respectively. ***: different from wildtype at P<0.001 (one-way ANOVA with Dunn’s post-hoc correction). ns: not significant.

We next determined the site of action of this gene by knocking out *gcy-12* in one or both AWC neurons. To do so, we obtained a *gcy-12* allele in which the first exon was flanked by *loxP* sites and expressed the Cre recombinase under the *srsx-3* promoter that drives expression strongly in AWC^OFF^ [32]. However, knocking out *gcy-12* in AWC^OFF^ had no effect on hexanol response plasticity such that these animals avoided hexanol similar to wild-type animals in sIAA (Fig 4B). In contrast, knocking out *gcy-12* in both AWC neurons by driving Cre expression under the bilateral *odr-1* promoter resulted in indifference to hexanol in sIAA similar to the phenotype of *gcy-12(null)* mutants (Fig 4B). Although it is possible that we were unable to achieve a complete knockout of *gcy-12* in AWC^OFF^, these results imply that GCY-12 function is required in both AWC neurons to regulate context-dependent hexanol response plasticity in AWC^OFF^.

### Mutations affecting left-right asymmetry in AWC abolish lateralization of hexanol response plasticity in *gcy-12* mutants

Mutations in the *nsy-1* MAPKKK and *nsy-5* innexin genes abolish AWC asymmetry, such that both neurons exhibit AWC^ON^ or AWC^OFF^ features in *nsy-1* and *nsy-5* mutants, respectively [29–31]. To ask whether these developmental pathways also affect context-dependent odor responses, we characterized *nsy-5* and *nsy-1* mutants as well as *nsy-5; gcy-12* and *gcy-12 nsy-1* double mutants. We reasoned that if mutations in *gcy-12* affect hexanol response plasticity only in AWC^OFF^, then *nsy-5; gcy-12* mutants with two AWC^OFF^ neurons would exhibit symmetric hexanol response plasticity defects in AWC. Conversely, neither AWC neuron is expected to show a hexanol response plasticity defect in *gcy-12 nsy-1* mutants with two AWC^ON^ neurons (see Fig 5H).

**Fig 5.**
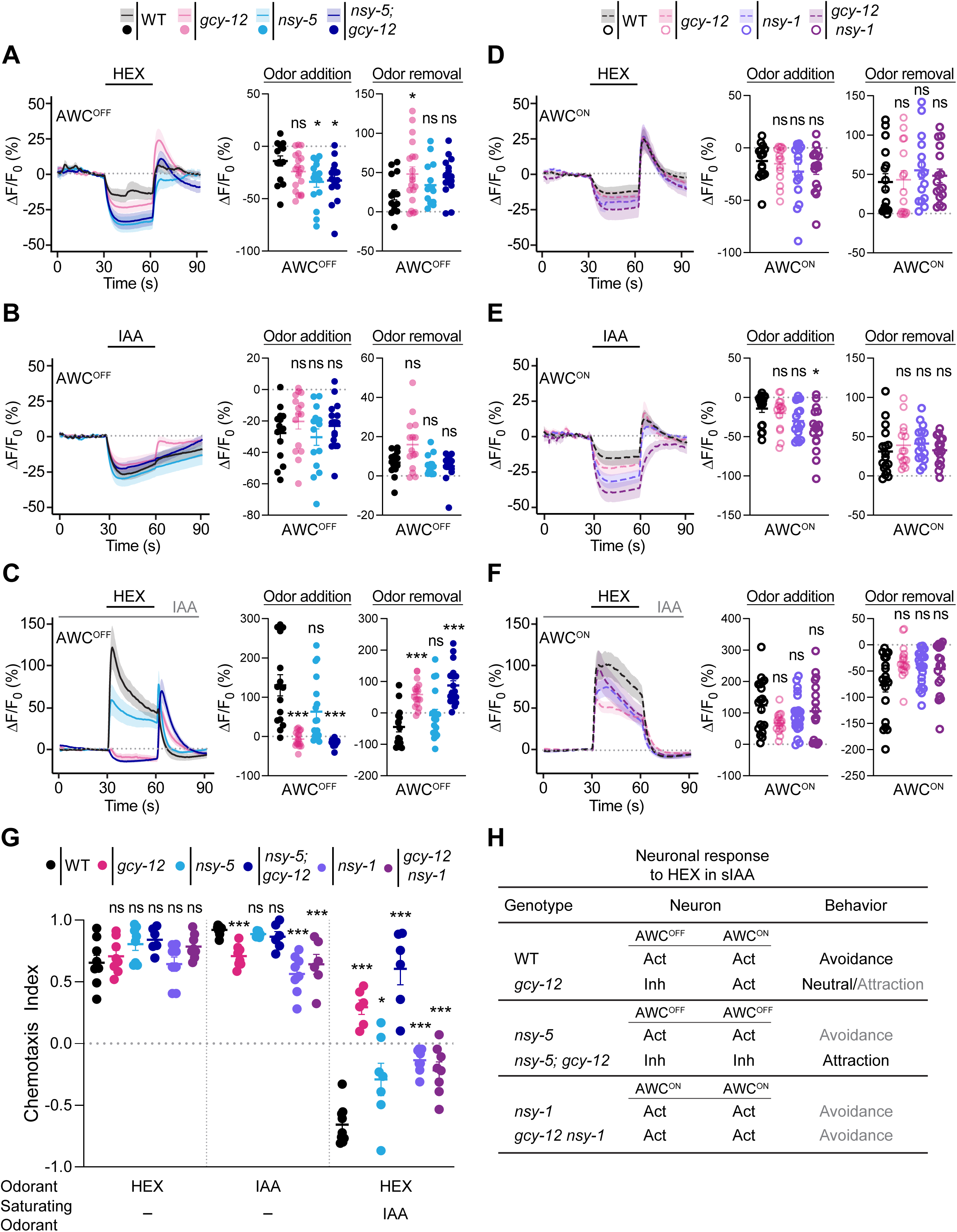
Loss of lateralization in AWC cell fate abolishes asymmetry in the response plasticity defects of *gcy-12* mutants. **A-F)** (Left) Average changes in GCaMP fluorescence in AWC^OFF^ and AWC^ON^ in the indicated genetic backgrounds upon addition of a 30 sec pulse of 10^-4^ hexanol (A,C,D,F) or IAA (B,E) indicated by a short solid line. The presence of a 10^-4^ concentration of sIAA in C and F is indicated by a long solid gray line. Shaded regions are SEM. (Right) Quantification of peak changes in fluorescence intensity in AWC^OFF^ or AWC^ON^ upon odorant addition or removal in each panel. Each circle is the value from a single neuron. Corresponding heatmaps are shown in S6 Fig. *gcy-12(ks100)* data are repeated from Fig 2 and indicated in light pink. Horizontal and vertical bars indicate the mean and SEM, respectively. Both AWC neurons are considered to be AWC^OFF^ in *nsy-5* and *nsy-5; gcy-12* mutants; both AWC neurons are considered to be AWC^ON^ in *nsy-1* and *gcy-12 nsy-1* mutants. * and ***: different from corresponding wildtype at P<0.05 and <0.001 (A: one-way ANOVA with Dunnett’s post-hoc correction; B-F: Kruskal-Wallis with Dunn’s post-hoc correction). ns: not significant. Alleles used were: *gcy-12(ks100), nsy-5(tm1896)*, and *nsy-1(ok390)*. **G)** Behavioral responses of animals containing the indicated mutations to point sources of hexanol with or without sIAA, and IAA alone. Concentrations of odorants used were 10^-1^ hexanol and 10^-2^ IAA as test odorants, and 10^-4^ IAA as the saturating odorant. Each dot is the chemotaxis index calculated from a single assay of ∼50-100 animals. Data shown are from at least three independent days. Horizontal and vertical bars are the mean and SEM, respectively. * and ***: different from corresponding wild-type at P<0.05 and 0.001, respectively (one-way ANOVA with Dunnett’s posthoc correction). ns: not significant. Alleles used were: *gcy-12(ks100), nsy-5(tm1896)*, and *nsy-1(ok390)*. **H)** Summary of expected and observed neuronal responses and behavioral phenotypes in animals of the indicated genotypes in response to hexanol in sIAA conditions. Act: Activated; Inh: Inhibited. Gray letters indicate observed weaker behavioral responses.

Addition of either hexanol or IAA in control conditions resulted in decreased intracellular calcium in both AWC neurons in wild-type, and all examined single and double mutants (Fig 5A, 5B, 5D, and 5E, S6A-D Fig). Hexanol response plasticity in sIAA was also unaffected in either AWC neuron in *nsy-1* and *nsy-5* single mutants (Fig 5C and 5F, S6E-F Fig). In sIAA, both neurons in *gcy-12 nsy-1* mutants were activated by hexanol consistent with both neurons exhibiting AWC^ON^ like properties (Fig 5F, S6F Fig). In contrast, both neurons in *nsy-5; gcy-12* mutants were inhibited by hexanol in sIAA, consistent with both neurons exhibiting AWC^OFF^ properties (Fig 5C, S6E Fig).

*nsy-1* mutants were previously shown to exhibit weak defects in attraction to IAA (Fig 5G) [28], and we found that both *nsy-1* and *nsy-5* single mutants also exhibited weaker avoidance of hexanol in sIAA possibly due to pleiotropic effects on other sensory neuron responses including those of ASH (Fig 5G) [31, 47, 48]. However, consistent with the symmetric hexanol response plasticity defects in *nsy-5; gcy-12* double mutants, these animals were strongly attracted to hexanol in sIAA (Fig 5G). Conversely, the behavioral phenotype of *gcy-12 nsy-1* double mutants remained similar to that of *nsy-1* mutants alone in response to hexanol in sIAA (Fig 5G), supporting the notion that mutations in *gcy-12* do not affect AWC^ON^ responses. These results (summarized in Fig 5H) confirm that GCY-12 regulates hexanol response plasticity asymmetrically in AWC^OFF^ neurons.

## DISCUSSION

Here we show that an olfactory neuron pair employs left-right asymmetric signaling pathways to exhibit symmetric context-dependent plasticity in its responses to a food-related odorant. Mutations in *gcy-12* affect hexanol response plasticity in sIAA only in AWC^OFF^ indicating that this plasticity in AWC^ON^ must be mediated in part via *gcy-12*-independent mechanisms. The contribution of distinct molecules and mechanisms to the sensory signal transduction cascade in AWC^OFF^ and AWC^ON^ under different conditions may increase the ability of these neurons to encode specific experiences in order to drive the appropriate adaptive behavior.

*C. elegans* moves towards or away from a point source of a chemical largely although not exclusively via modulation of its turning frequency [49, 50]. Inhibition and activation of both AWC neurons upon odorant addition or removal decreases and increases turns, respectively, thereby driving animals up an attractive odor gradient [38, 45, 51]. In sIAA, addition and removal of hexanol instead activate and inhibit both AWC neurons, resulting in avoidance of this chemical. However, in *gcy-12* mutants, AWC^OFF^ and AWC^ON^ exhibit conflicting responses to hexanol in sIAA, with AWC^ON^ being strongly activated upon hexanol addition, and AWC^OFF^ being strongly inhibited upon hexanol removal. Thus, while responses in AWC^ON^ are expected to drive hexanol avoidance, AWC^OFF^ responses promote attraction. Consequently, animals are largely indifferent to hexanol under these conditions. This defect in behavioral plasticity is distinct from the symmetric response defect of *odr-3* mutants or those of *nsy-5; gcy-12* double mutants, in which both AWC neurons remain inhibited by hexanol in sIAA [37], resulting in strong attraction to this odorant.

The response profiles of AWC^ON^ and AWC^OFF^ to odorants are partly distinct, and each AWC neuron expresses a partly distinct set of chemoreceptors [28, 29, 32, 33, 38, 52, 53]. While each AWC neuron also expresses multiple members of additional sensory signaling protein families [23, 54, 55], the expression of many sensory signaling molecules is largely symmetric. In principle, the asymmetric expression of chemoreceptors alone may be sufficient to lateralize AWC responses. However, our results indicate that under distinct contexts, chemoreceptors may engage with different signaling pathways in AWC^ON^ and AWC^OFF^ to mediate response plasticity. Since neither the hexanol nor the IAA receptors in AWC have yet been identified, it is currently unclear whether distinct chemoreceptors mediate responses to these odorants in each AWC neuron under different conditions. Given the strong effect of *gcy-12* mutations specifically on hexanol response plasticity in AWC^OFF^, it is possible that chemoreceptors for other odorants such as heptanol couple with alternative signaling pathways symmetrically or asymmetrically to modulate behavior. The recruitment of distinct signaling molecules in different contexts may account for the co-expression of multiple members of gene families encoding signal transduction proteins in individual sensory neurons in *C. elegans*.

Functional asymmetry in sensory neuron response properties can arise via cell autonomous or cell non-autonomous mechanisms. We previously showed that the switch in the sign of the hexanol response in sIAA in AWC is independent of chemical or neuropeptidergic transmission, suggesting that this response plasticity may be mediated cell-autonomously [37]. However, our finding that *gcy-12* function must be lost in both AWC neurons to phenocopy the behavioral defect of *gcy-12(null)* mutants leads us to speculate that these neurons may in addition communicate to modulate responses cell non-autonomously. Thus, in the absence of *gcy-12* function only in AWC^OFF^, plasticity in AWC^ON^ may be sufficient to drive response plasticity in AWC^OFF^, thereby promoting robust hexanol avoidance. In this model, loss of *gcy-12* in both AWC neurons affects response plasticity in AWC^OFF^ via both cell autonomous and cell non-autonomous mechanisms without affecting AWC^ON^ responses, thereby resulting in indifference to hexanol. We note that while ablating only AWC^OFF^ or AWC^ON^ is sufficient to abolish attraction to the odorants 2,3-pentanedione or butanone, respectively [28], both AWC neurons respond to these odorants [53] (A. Pandey, unpublished results), raising the possibility that responses in one neuron may drive responses in their bilateral partner. Although no gap junctions between the two AWC neurons have been reported in postembryonic animals [56, 57], these neurons may communicate via a small number of chemical synapses (www.wormwiring.org) [57]. Whether and how the two AWC neurons communicate, and a possible role of GCY-12 in this communication, remain to be determined.

Lateralization of sensory neuron responses and behavior may be mediated via asymmetric expression of signaling proteins, isoform usage, trafficking mechanisms, synaptic connectivity, neurotransmitter release, and neuron number variation among others (eg. [18, 25, 29, 42, 52, 58–60]). In addition to genetically hard-wired mechanisms, experience can modulate neuronal asymmetry. For instance, the lateralization of synaptic connectivity of ASEL and ASER is switched as a function of experience [58], honeybees and *Drosophila* exhibit time-dependent lateralization in olfactory learning behaviors [61–63], and asymmetric light stimulation in early development regulates lateralization of the visual system in chickens and pigeons [64, 65]. Our observations suggest that lateralization in neuronal properties may be revealed only in specific genetic backgrounds or environmental contexts, suggesting that functional and mechanistic asymmetry in seemingly bilaterally symmetric neurons may be a more general feature of neuronal responses. We propose that this asymmetry further diversifies sensory neuron functions, particularly in animals with polymodal sensory neurons, and enables specific context-dependent behavioral flexibility.

## MATERIALS and METHODS

### *C. elegans* growth and genetics

The wild-type strain was *C. elegans* Bristol N2. Worms were maintained at 20°C on nematode growth medium (NGM) plates seeded with *E. coli* OP50. All experiments were performed using well-fed one-day old adult hermaphrodites. Animals were grown under well-fed conditions for at least two generations prior to testing. The presence of mutations was verified by PCR and/or Sanger sequencing. Strains used in this study are listed in S1 Table.

### Generation of gene-edited alleles

CRISPR alleles were isolated as described [66]. Cas9 protein and guide RNAs were ordered from IDT. Repair templates were PCR amplified from the relevant plasmid and melted before adding to the injection mix. The injection mix included the pRF4 *rol-6(gf)* plasmid. Injected animals were placed on individual plates, and 96 F1 progeny were singled from plates that contained rollers, allowed to reproduce until the plate was starved, and were then screened by PCR for the expected change. Insertion of *gfp* sequences to generate a *gcy-12.a* reporter tagged allele was performed first, and *mScarlet* sequences were subsequently inserted into this allele to also tag *gcy-12.b*.

The floxed *gcy-12* allele (*gcy-12(syb10363)* was generated by Suny Biotech. LoxP sites were inserted 283 bp apart flanking the first exon. The *gcy-12(oy213)* allele was generated by replacing C with T (Q562 to STOP) via a repair template from IDT in a *nsy-1(ok390)* containing background to obtain the PY12512 strain. The amino acid change in *gcy-12(oy213)* is identical to that in *gcy-12(ks100)*.

### Molecular biology

A 1068 bp *odr-1* promoter or a 1327 bp *srsx-3* promoter were used to drive *Cre::SL2::BFP* or *Cre::SL2::GFP.* NLS sequences were fused to both Cre and BFP/GFP. *odr-1*p::*Cre::SL2::GFP* was injected at 10 ng/μl with the *unc-122*p*::gfp* co-injection marker at 50 ng/μl. Remaining Cre-expressing plasmids were injected at 25 ng/µl with a co-injection marker (*vha-6*p*::NLS-gfp*) and DNA ladder (NEB) injected at 10 ng/µl and 65 ng/µl, respectively. To improve CRE efficiency, the SV40 NLS fused to Cre was replaced with the NLS from *egl-13* [67, 68]. This optimized Cre (optCre) was injected at 50 ng/µl with the co-injection marker (*vha-6*p*::NLS-gfp*) and DNA ladder (NEB) injected at 10 ng/µl and 40 ng/µl, respectively.

For *gcy-12* splice isoform localization, a 1068 bp *odr-1* promoter was used to express *gcy-12.a::mStayGold* or *gcy-12.b::mScarlet-13*. Both plasmids were injected together at 30 ng/µl and a co-injection marker (*vha-6*p*::NLS-gfp*) and DNA ladder (NEB) at 10 ng/µl and 30ng/µl, respectively. Plasmids used in this work are listed in S2 Table.

### Behavioral assays

Chemotaxis behavioral assays were performed as previously described [20, 37, 69]. Assays were performed on 10 cm square plates with the exception of attraction assays to IAA or benzaldehyde alone which were performed on 10 cm round plates. For saturation assays, the relevant odorant was added to the agar prior to pouring plates (1 μl odorant at 10^-4^ dilution/10 ml of assay agar). Odorants were freshly diluted in ethanol each day. Unless indicated otherwise, 1 μl of diluted odorant or ethanol was placed at each of two spots on either side of a square assay plate, or at one spot on round assay plates. 1 μl of 1M sodium azide was added to each odorant or diluent spot to paralyze animals.

10-15 L4 larvae were picked onto 10 cm plates seeded with 1 ml of OP50 four days prior to the behavioral assay. On the day of the assay, animals were washed off growth plates with S-basal buffer, washed two more times with S-basal buffer, and once with milliQ water. 50-200 washed worms were placed at the center of the assay plate and allowed to move for an hour. The chemotaxis index was calculated as: [(animals in the two horizontal rows adjacent to the odor (square plates) or animals at the odor (round plates)] – [(animals in the two rows adjacent to the diluent (square plates) or animals at the diluent (round plates)]/total number of animals. For strains carrying extrachromosomal arrays, only animals expressing the fluorescent co-injection marker were included in the quantification. 2-3 assay plates were tested each day per genotype and condition; data reported are from biologically independent experiments from at least 3 days.

### Calcium imaging

Calcium imaging was performed using modified two-layer microfluidics devices [37, 70]. All buffers and odorant dilutions were prepared the day of the experiment. Test odorants were diluted to 10^-4^ in S-basal which was also used as the control buffer. For saturation experiments, all channels in the microfluidics device contained the saturating odorant diluted to 10^-4^ in S-basal. 1 μl of 20 μM fluorescein was used to visualize buffer flow. Both AWC neurons respond robustly to shear stress as a consequence of fluid flow in the microfluidics device [71, 72]. Shear stress generated due to rapid fluid flow in the imaging devices can result in altered and variable AWC responses to test odorants. We previously performed imaging in the presence of buffer alone to ensure that fluid flow did not evoke responses in AWC under the conditions used [37]. We also interspersed wild-type and mutant animals in each imaging session to minimize artefactual responses. Imaging sessions in which wild-type AWC neurons responded variably due to fluctuations in fluid flow in the system were terminated. In future, imaging of odorant-evoked stimuli in AWC may benefit from the use of shearless microfluidics devices [71].

Prior to imaging, one-day old adults were picked onto an unseeded NGM plate and allowed to immobilize in the loading buffer containing poloxamer 188 (Sigma), 1 mM tetramisole hydrochloride, and the saturating odorant diluted in S-basal for 10 min before loading into the microfluidics device. Imaging was performed for 1 cycle of 30 sec buffer/30 sec odor/30 sec buffer on an Olympus BX52WI microscope with a 40x oil objective. Images were captured with a Hamamatsu Orca CCD camera for 90 secs at 4 frames per second. The exposure was set to 250 milliseconds with 4×4 binning.

GCaMP fluorescence intensity changes were quantified using the register_quantGCaMP.ijm macro in Fiji ImageJ (https://imagej.net/software/fiji/). The region of interest (ROI) was manually outlined around the soma or cilia. Background fluorescence intensity was subtracted from the ROI and this value was used for further analysis. Data were visualized and figures were generated using RStudio (version 4.2.764). F_0_ was determined as the average fluorescence in the 5 secs prior to odor onset. To correct for photobleaching, an exponential function was fit to the fluorescence intensity values for the first and last 20 secs of an imaging session, and the curve was subtracted from the raw ΔF/F_0_ values at each time point. Peak amplitudes for odor addition and removal were quantified as the maximum amplitude change (F-F_0_) in the first 10 secs of stimulus addition or removal. Custom scripts for analyses of calcium imaging data are available at https://doi.org/10.5281/zenodo.15939087. AWC^OFF^ and AWC^ON^ neurons were distinguished via expression of *srsx-3*p*::mScarlet* in AWC^OFF^ (see S1 Table). Wild-type and mutant animals were interspersed throughout each imaging session. 5-10 animals per genotype and condition were examined in each session; data reported are from at least two biologically independent experiments.

### Imaging of fluorescent reporter expression

Animals were anaesthetized with 100 mM levamisole (Sigma Aldrich) on 10% agarose pads on microscope slides. Transgenic animals expressing *gcy-12a::gfp* and *gcy-12b::mScarlet* were imaged on a Zeiss LSM880 AiryScan confocal microscope using a 63x oil immersion objective. Transgenic animals expressing reporter-tagged *gcy-12* isoforms were imaged using a 40x oil immersion objective on an inverted spinning disk confocal microscope (Zeiss Axiovert with a Yokogawa CSU22 spinning disk confocal head and a Photometrics Quantum SC 512 camera) and Slidebook 6.0 (Intelligent Imaging Innovations, 3i) software.

### Statistical analyses

Behavioral and calcium imaging data were plotted and analyzed in GraphPad Prism v10.5 (www.graphpad.com). All datasets were first tested for normality via a Shapiro-Wilk test with a P-value set at 0.05. For normal data, datasets with two samples were compared using an unpaired t-test with a post-hoc Welch’s correction, and datasets with more than two samples were compared using one-way ANOVA and a post-hoc Dunnett’s multiple comparisons correction. For non-normal data, datasets with two samples were compared using a Mann-Whitney-Wilcoxon t-test, and datasets with more than two samples were compared using a Kruskal-Wallis test and a post-hoc Dunn’s multiple comparisons correction. Statistical tests used are indicated in each figure legend.

## Acknowledgements

We are grateful to Michael O’Donnell, Munzareen Khan, Kirsten Judge, Priya Dutta and Jihye Yeon for advice and technical assistance, Frank Mello for assistance with troubleshooting calcium imaging conditions, Jinmahn Kim (Northeastern University) for assistance with generating microfluidics devices for calcium imaging, the *Caenorhabditis* Genetics Center for strains, and the Brandeis Materials Research Science and Engineering Center (MRSEC) for access to the microfabrication facility. We thank Cori Bargmann, Anna Hartmann, Munzareen Khan, Michael O’Donnell, and members of the Sengupta lab for advice and critical comments on the manuscript. This work was funded in part by the NSF (IOS 2042100 – P.S.) and the NIH (R35 GM122463 – P.S.).

## Author Contributions

Conceptualization: A. Pandey, P. Sengupta.; Methodology: A. Pandey, M. Katz, S. Nurrish, A. Philbrook; Investigation: A. Pandey, M. Katz, S. Nurrish, A. Philbrook; Writing - original draft: A. Pandey, P. Sengupta; Writing - review and editing: A. Pandey, M. Katz, S. Nurrish, A. Philbrook, P. Sengupta; Visualization: A. Pandey, A. Philbrook; Supervision: P. Sengupta; Funding acquisition: P. Sengupta.

## S1 Data

Numerical data for all behavioral and calcium imaging experiments are provided at https://doi.org/10.5281/zenodo.15939087.

## SUPPLEMENTARY FIGURE LEGENDS

**Fig S1.**
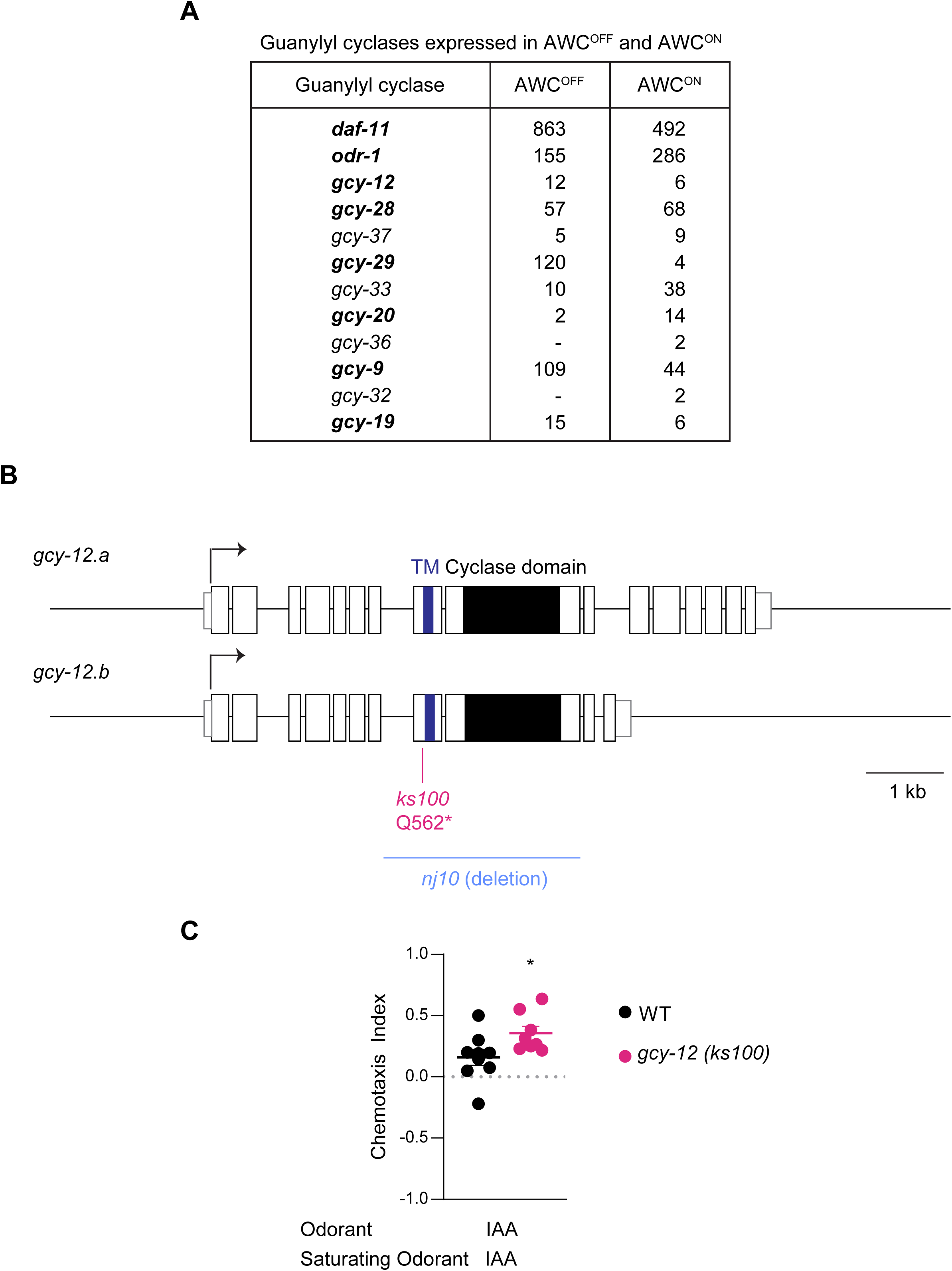
AWC-expressed guanylyl cyclase genes. **A)** Expression levels of indicated receptor and soluble guanylyl cyclase genes in the two AWC neurons from CeNGEN [33]. Expression values are in transcripts per million (TPM) without thresholding (www.cengen.org) [33]. Receptor guanylyl cyclases are bolded. **B)** Gene structure of *gcy-12* and the encoded *gcy-12.a* and *gcy-12.b* transcript isoforms. The molecular lesions in the *ks100* and *nj10* alleles are indicated [43]. **C)** Behavioral responses of wild-type and *gcy-12(ks100)* animals to a point source of 10^-3^ IAA in 10^-4^ IAA as the saturating odorant. Each dot is the chemotaxis index calculated from a single assay of ∼50-100 animals. Data shown are from at least three independent days. Horizontal and vertical bars are the mean and SEM, respectively. *: different from wild-type at P<0.05 (unpaired t-test with a post-hoc Welch’s correction).

**S2 Fig.**
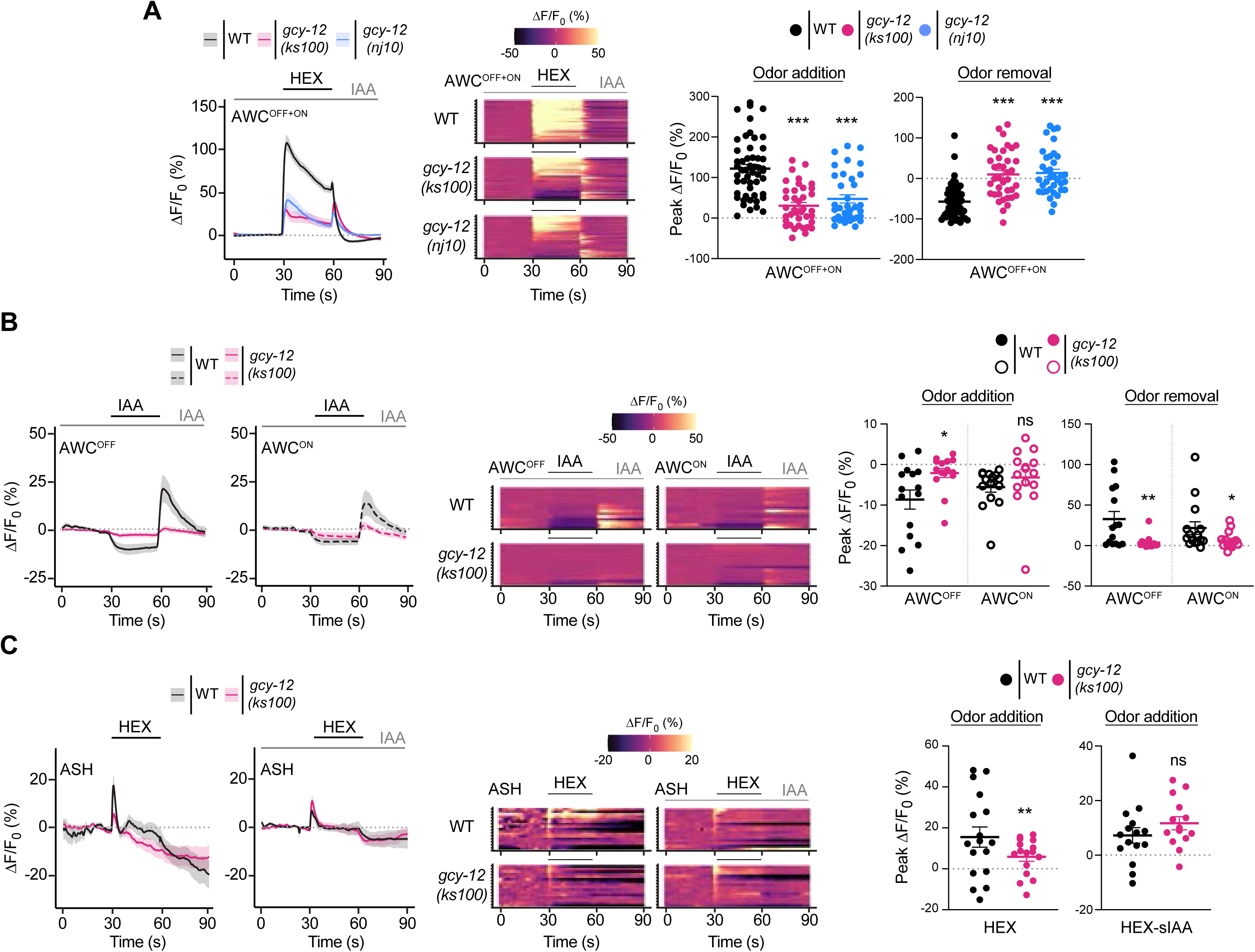
Mutations in *gcy-12* affect context-dependent hexanol response plasticity only in AWC^OFF^. **A-C)** (Left) Average changes in GCaMP fluorescence in AWC (A, B) or ASH (C) neurons in wild-type and *gcy-12* mutants upon addition of a 30 sec pulse of 10^-4^ hexanol (A, C) or IAA (B) indicated by a short solid line in 10^-4^ concentration of sIAA (long solid gray line). Shaded regions are SEM. (Middle) Heatmaps of changes in fluorescence intensity corresponding to the responses shown at left in each panel. Each row in the heatmaps shows responses from a single neuron. (Right) Quantification of peak changes in fluorescence intensity upon odorant addition and/or removal in each panel. Each circle is the value from a single neuron. Horizontal and vertical bars indicate the mean and SEM, respectively. *, **, and ***: different from corresponding wild-type at P<0.05, 0.01 and 0.001, respectively (A: Kruskal-Wallis with Dunn’s post-hoc correction; B, C: Mann-Whitney-Wilcoxon test); ns: not significant. Data in A are repeated in Fig 2C, separated by AWC^OFF^ and AWC^ON^ neurons.

**S3 Fig.**
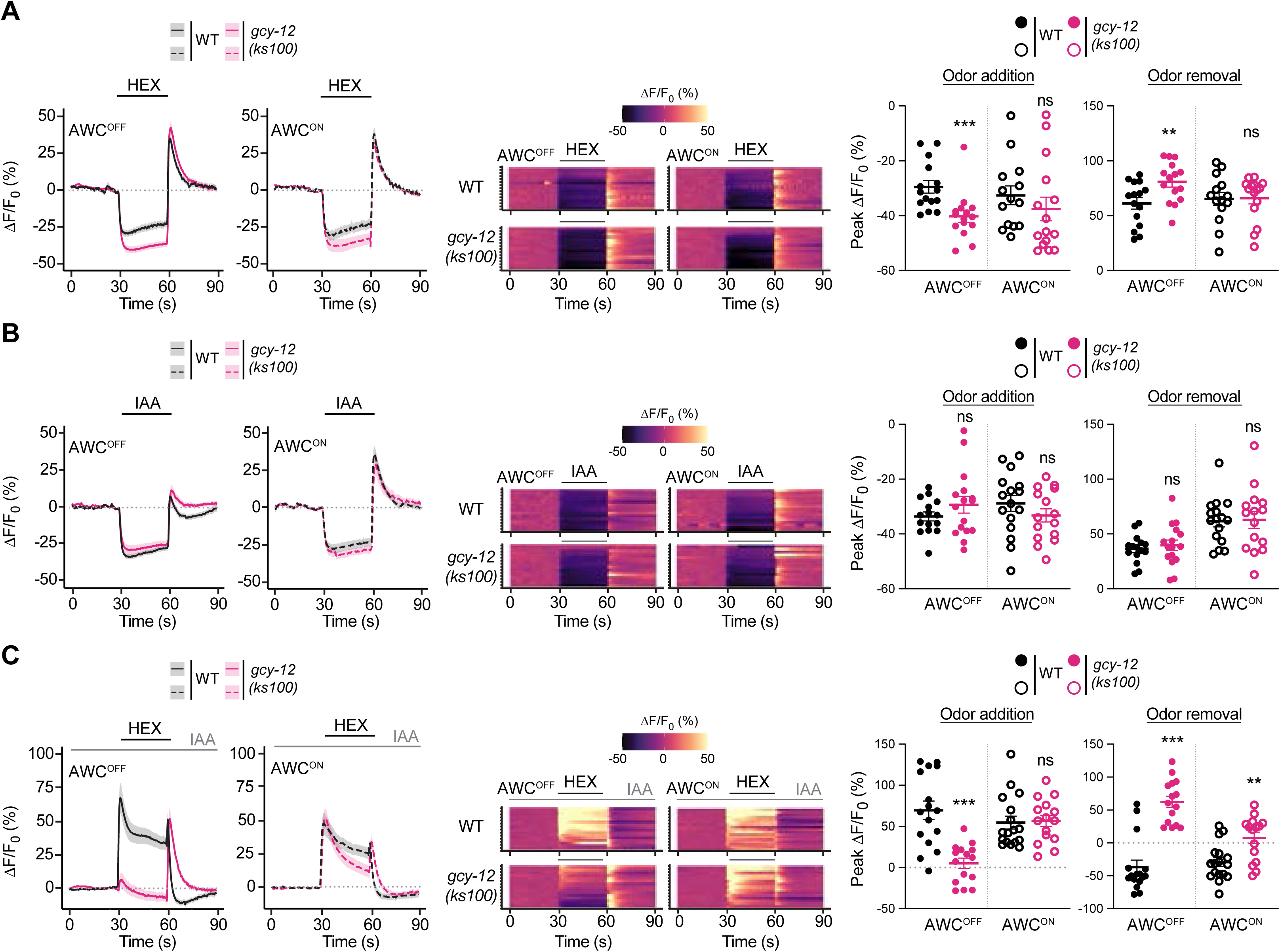
*gcy-12* mutants exhibit asymmetric hexanol-evoked calcium responses in sIAA in AWC neuronal cilia. **A-C)** (Left) Average changes in GCaMP fluorescence in the cilia of AWC neurons in wild-type and *gcy-12* mutants upon addition of a 30 sec pulse of 10^-4^ hexanol (A) or IAA (B) indicated by a short solid line, and 10^-4^ hexanol in 10^-4^ concentration of sIAA (C, long solid gray line). Shaded regions are SEM. (Middle) Heatmaps of changes in fluorescence intensity corresponding to the responses shown at left in each panel. Each row in the heatmaps shows responses from a single neuron. (Right) Quantification of peak changes in fluorescence intensity upon odorant addition or removal in each panel. Each circle is the value from a single neuron. Horizontal and vertical bars indicate the mean and SEM, respectively. ** and ***: different from corresponding wild-type at P<0.01 and 0.001, respectively (A, C: Mann-Whitney-Wilcoxon test, B: unpaired t-test with a post-hoc Welch’s correction n); ns: not significant.

**S4 Fig.**
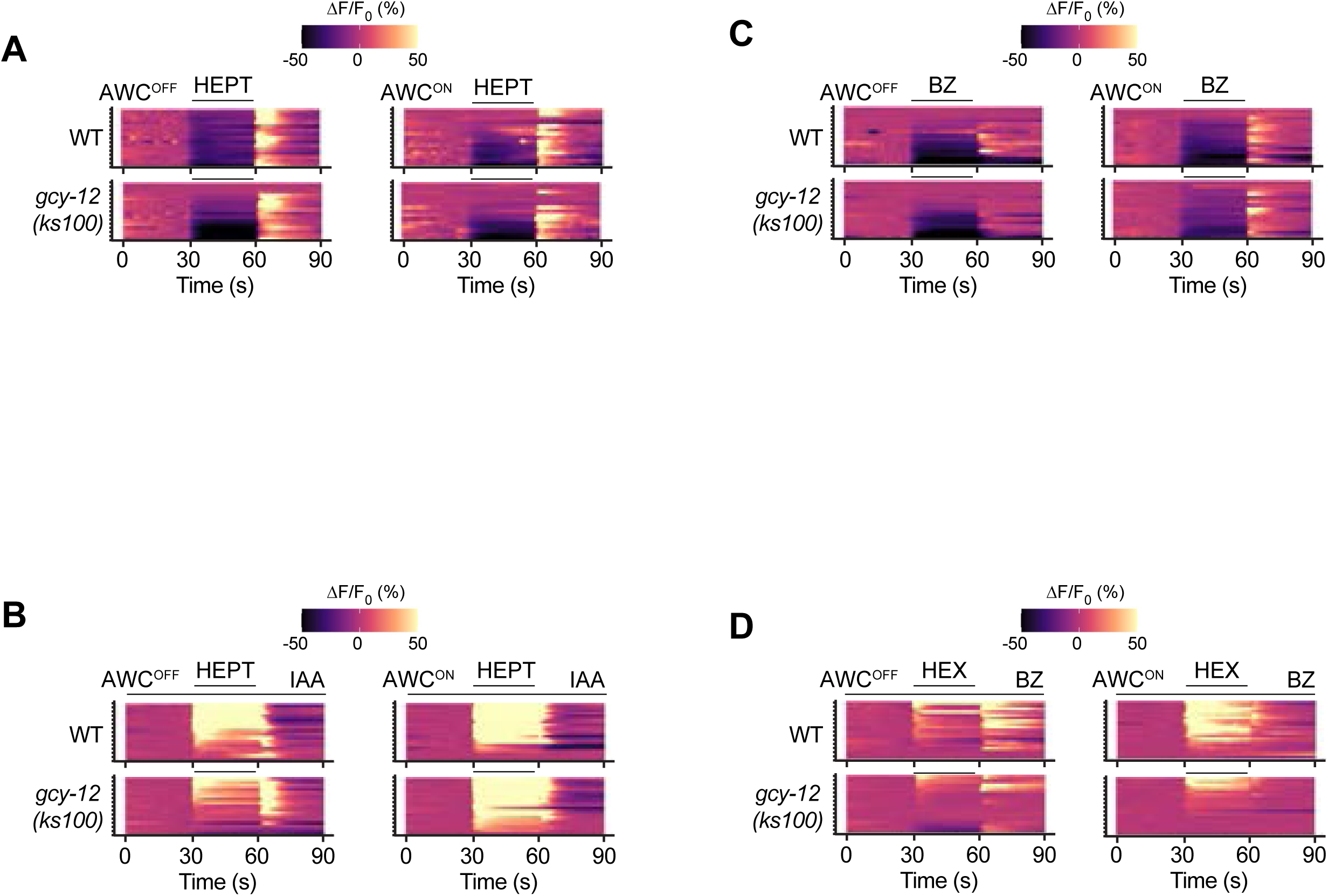
Context-dependent plasticity defects in *gcy-12* mutants are partly odorant-specific. **A-D)** Heatmaps of changes in fluorescence intensity corresponding to the responses shown in Fig 3B, 3C, 3E and 3F. Each row in the heatmaps shows responses from a single AWC^OFF^ or AWC^ON^ neuron from different animals.

**S5 Fig.**
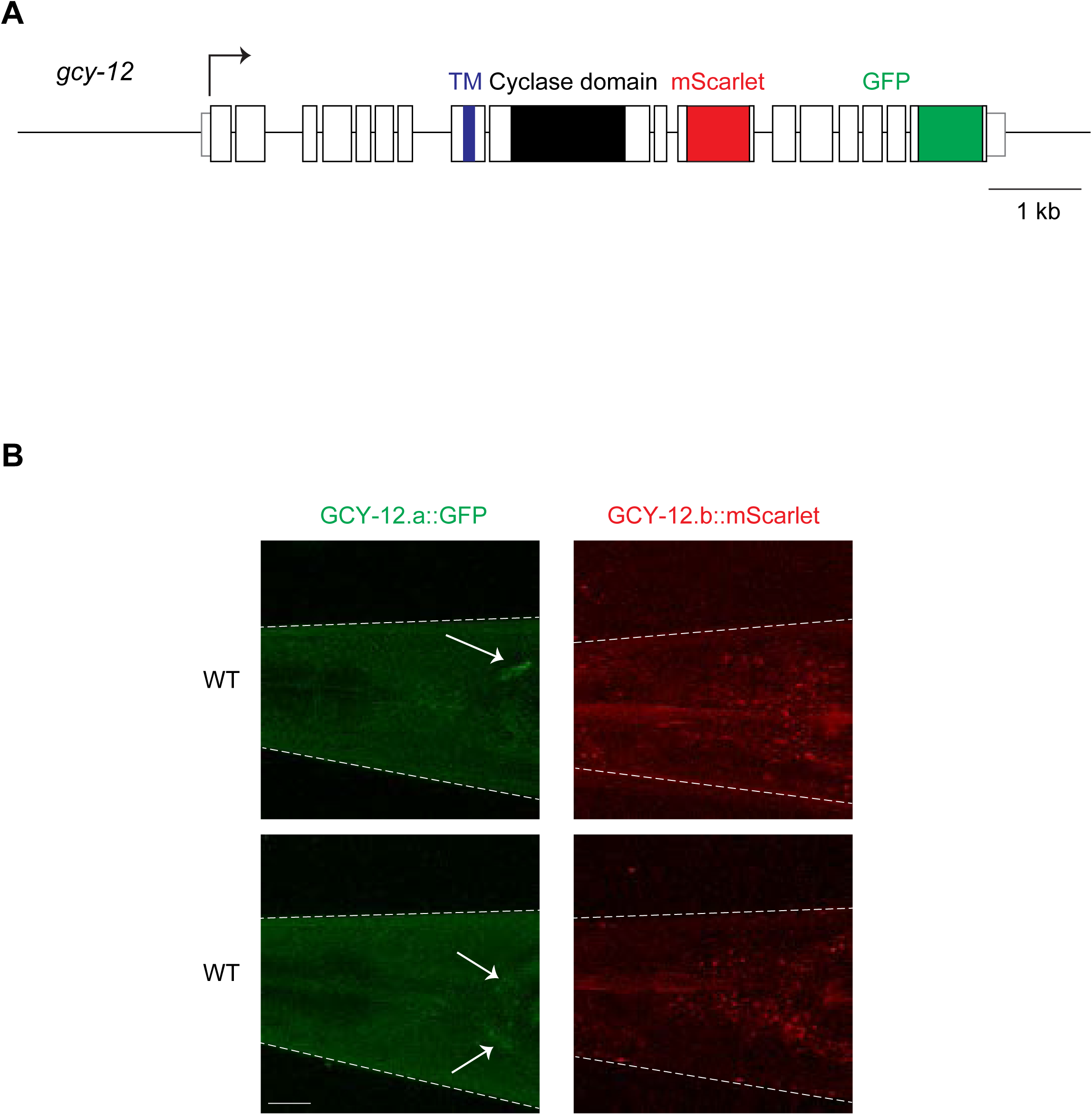
Loss of *gcy-12* only in AWC^OFF^ is not sufficient to alter hexanol response plasticity in sIAA. **A)** Genomic structure of *gcy-12* indicating the sites of insertion of fluorescent reporter sequences at the endogenous locus. Also see S1B Fig. **B)** Representative images showing expression of the two reporter-tagged *gcy-12* isoforms in the head. Arrows indicate putative neuronal soma. Expression of *gcy-12.b* was undetectable. Anterior at left. Scale bar: 5 μm.

**S6 Fig.**
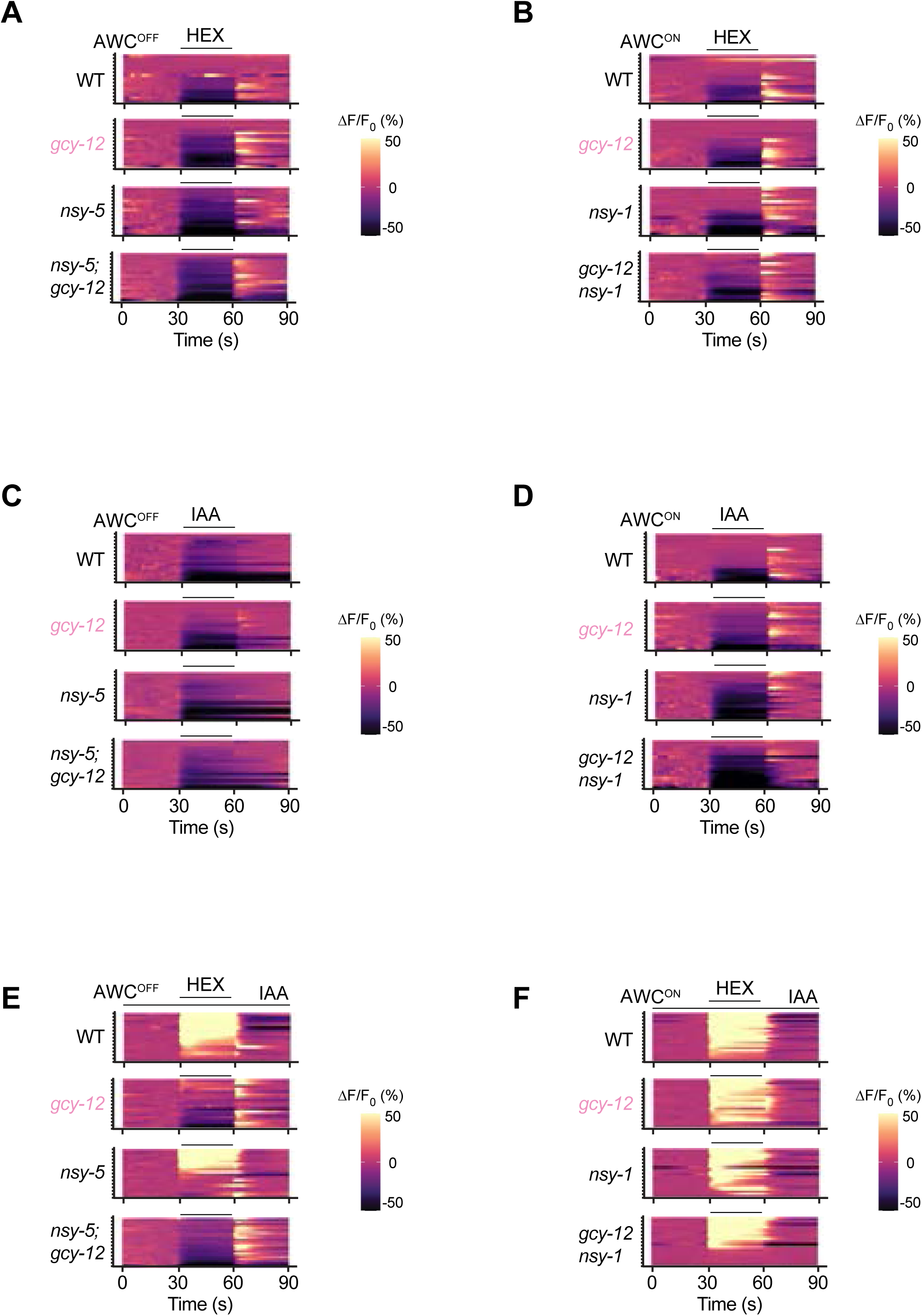
Loss of AWC fate asymmetry abolishes asymmetry in the odorant response defects of *gcy-12* mutants. **A-F)** Heatmaps of changes in fluorescence intensity corresponding to the responses shown in Fig 5A-F. Each row in the heatmaps shows responses from a single AWC^OFF^ or AWC^ON^ neuron from different animals. Both AWC neurons are considered to be AWC^OFF^ in *nsy-5* and *nsy-5; gcy-12* mutants; both AWC neurons are considered to be AWC^ON^ in *nsy-1* and *gcy-12 nsy-1* mutants. *gcy-12* data are repeated from Fig 2 and indicated in light pink.

**S1 Video.** Intracellular calcium levels in AWC^ON^ are increased upon addition of hexanol in both wild-type and *gcy-12* mutants.

Fluorescence changes in AWC^ON^ in wild-type (left) and *gcy-12(ks100)* (right) animals expressing GCaMP3 in response to a 30 sec pulse of 10^-4^ hexanol in 10^-4^ sIAA. Video is at 6X speed.

**S2 Video.** Intracellular calcium levels are increased in AWC^OFF^ upon addition or removal of hexanol in wild-type and *gcy-12* mutants, respectively.

Fluorescence changes in AWC^OFF^ in wild-type (left) and *gcy-12(ks100)* (right) animals in response to a 30 sec pulse of 10^-4^ hexanol in 10^-4^ sIAA. Video is at 6X speed.

**S1 Table.**
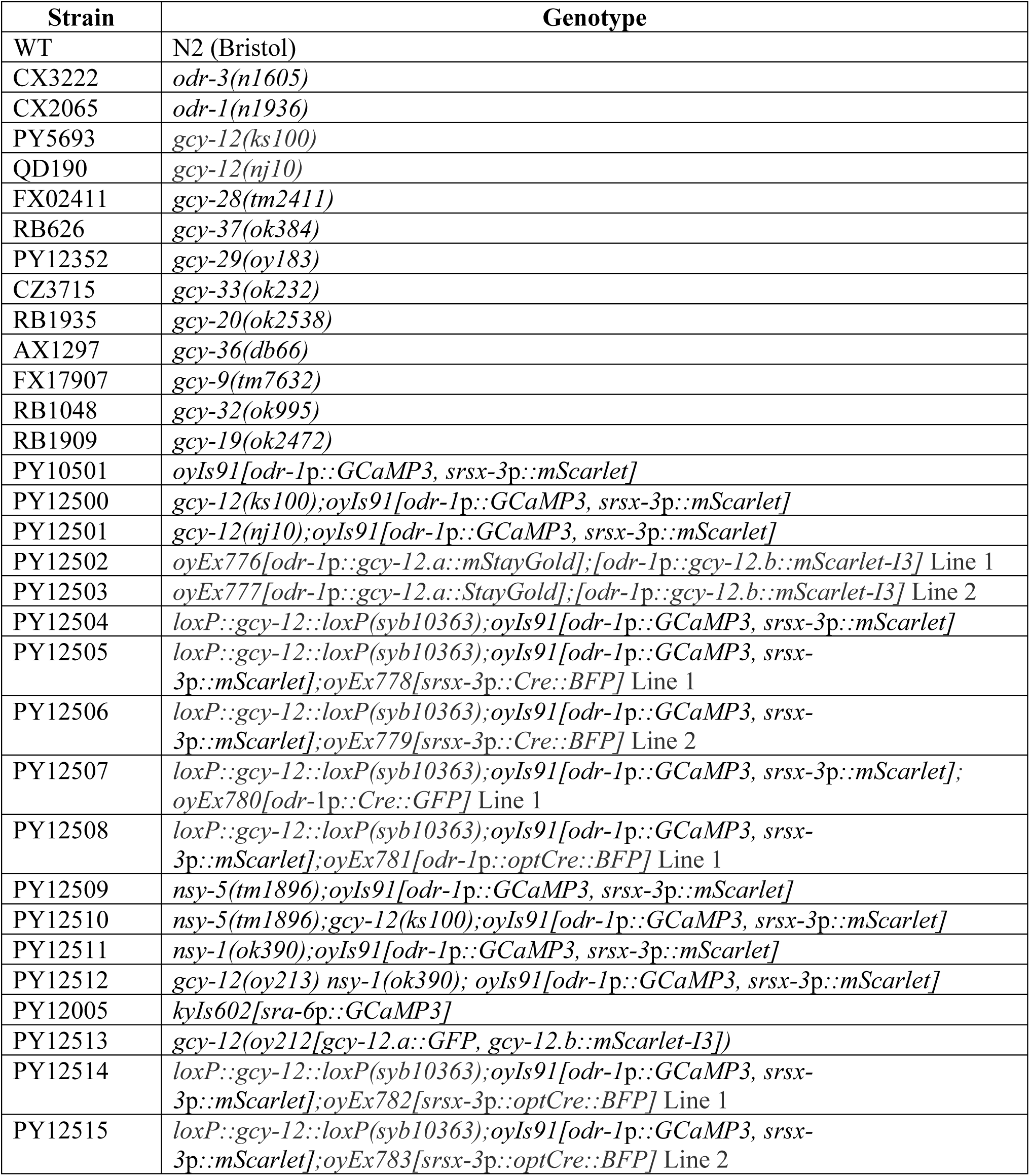
Strains used in this work.

**S2 Table.** Plasmids used in this work.

| Plasmid | Description |
| --- | --- |
| PSAB1387 | *odr-1p::gcy-12.a::mStayGold* |
| PSAB1388 | *odr-1p::gcy-12.b::mScarlet-I3* |
| PSAB1389 | *odr-1p::SV40NLS-Cre::SL2::NLS-GFP* |
| PSAB1390 | *srsx-3p::SV40NLS-Cre::SL2::NLS-BFP* |
| PSAB1391 | *odr-1p::egl-13NLS-Cre::SL2::NLS-BFP* |
| PSAB1392 | *srsx-3p::egl-13NLS-CRE::SL2::NLS-BFP* |

